# DTPSP: A Deep Learning Framework for Optimized Time Point Selection in Time-Series Single-Cell Studies

**DOI:** 10.1101/2024.12.18.629276

**Authors:** Michel Hijazin, Pumeng Shi, Jingtao Wang, Jun Ding

**Author notes:** These authors contributed equally to this work (Michel Hijazin, Pumeng Shi, Jingtao Wang).

## Abstract

Time-series studies are critical for uncovering dynamic biological processes, but achieving comprehensive profiling and resolution across multiple time points and modalities (multi-omics) remains challenging due to cost and scalability constraints. Current methods for studying temporal dynamics, whether at the bulk or single-cell level, often require extensive sampling, making it impractical to deeply profile all time points and modalities. To overcome these limitations, we present DTPSP, a deep learning framework designed to identify the most informative time points in any time-series study, enabling resource-efficient and targeted analyses. DTPSP models temporal gene expression patterns using readily obtainable data, such as bulk RNA-seq, to select time points that capture key system dynamics. It also integrates a deep generative module to infer data for non-sampled time points based on the selected time points, reconstructing the full temporal trajectory. This dual capability enables DTPSP to prioritize key time points for in-depth profiling, such as single-cell sequencing or multi-omics analyses, while filling gaps in the temporal landscape with high fidelity. We apply DTPSP to developmental and disease-associated time courses, demonstrating its ability to optimize experimental designs across bulk and single-cell studies. By reducing costs, enabling strategic multi-omics profiling, and enhancing biological insights, DTPSP provides a scalable and generalized solution for investigating dynamic systems.

## Introduction

Time-series experiments are indispensable for uncovering dynamic biological processes, offering crucial insights into how cellular and molecular states evolve over time. These studies have been pivotal in developmental biology [1, 2], stem cell differentiation [3, 4], immune responses [5], and stress responses [6]. While bulk RNA sequencing has historically dominated time-series analyses due to its affordability, it averages gene expression across thousands of cells, obscuring rare or transient states critical for understanding biological heterogeneity. By contrast, single-cell profiling [7] captures these nuances, revealing rare cell populations and subtle transitions with unprecedented resolution. Similarly, multiomics approaches integrate transcriptomics, proteomics, and epigenomics, offering a comprehensive view of cellular states and uncovering complex regulatory networks. Despite these advancements, profiling multiple time points in single-cell or multi-omics studies remains prohibitively expensive, rendering comprehensive temporal profiling both financially and logistically challenging. Moreover, not all time points are equally informative; redundant or uninformative sampling inflates costs while yielding limited biological value. Identifying the most informative time points is therefore a critical challenge in time-series single-cell and multiomics studies.

Existing methods for time-point selection face notable limitations, particularly in high-throughput, high-resolution studies. Traditional approaches, such as uniform sampling intervals [8] and phenotype-driven heuristic strategies [9, 10], are simple but often fail to capture key temporal transitions in complex biological systems. Iterative methods, including those by Singh et al. [11] and Rosa et al. [12], dynamically refine time point selection using previously collected data. While these approaches improve temporal resolution, they introduce batch effects, increase experimental complexity, and rely heavily on prior knowledge. The Time Point Selection (TPS) method [13], which uses spline interpolation [14] to minimize prediction error, offers another alternative. However, TPS has significant shortcomings: it neglects gene interdependencies, has limited predictive power, and is ill-suited for the high-dimensional nature of single-cell data. Furthermore, current methods cannot predict or generate data for unobserved time points from bulk observations—a critical limitation in single-cell studies where exhaustive sampling is infeasible. As a result, these methods struggle to model dynamic biological changes effectively, reducing their utility in longitudinal studies.

To address these challenges, we introduce the Deep Time Point Selector and Profiler (DTPSP), a deep learning-based framework [15] specifically designed to optimize time-point selection for cost-effective, high-resolution time-series studies. DTPSP identifies the most informative time points while minimizing redundancy, ensuring maximal temporal resolution in single-cell and multiomics studies. Beyond time-point selection, DTPSP leverages predictive capabilities to reconstruct gene expression trajectories across unobserved time points using data from a limited set of sampled points. By integrating the deep generative module adapted from scSemiProfiler [16], DTPSP further enables the generation of single-cell-level trajectories for unprofiled time points, empowering researchers to perform comprehensive single-cell analyses without exhaustive profiling. Unlike existing approaches, DTPSP combines efficiency and accuracy, offering a scalable solution that overcomes the financial and logistical barriers of high-resolution time-series studies.

We validated DTPSP on datasets from three time-series transcriptomics studies, demonstrating its ability to accurately reconstruct single-cell trajectories that align closely with real sequencing data. These reconstructed trajectories enable robust downstream analyses, producing results comparable to those obtained from actual single-cell profiling. Additionally, DTPSP-selected time points are much more informative than time points selected via clustering analysis, achieving lower single-cell time-series trajectory reconstruction errors.

DTPSP addresses the prohibitive costs and experimental complexities associated with comprehensive time-series single-cell and multi-omics studies by offering a cost-effective and efficient strategy for selecting and profiling critical time points. This approach enables longitudinal studies that were previously impractical, allowing researchers to track dynamic biological processes with precision. For example, DTPSP can facilitate the mapping of differentiation trajectories in developmental biology [17, 18], the longitudinal monitoring of immune responses in immunology [19, 20], cancer research [21], aging and degenerative diseases [22], and the identification of key cellular transitions during tissue repair in regenerative medicine [23, 24]. By reducing experimental redundancy and focusing on the most informative time points, DTPSP enhances the ability to uncover temporal mechanisms critical to understanding complex biological systems while minimizing resource use.

## Results

### Methods overview

DTPSP is designed to optimize time-series studies by leveraging more accessible and cost-effective profiling, such as bulk RNA-seq, to identify the most informative time points for further in-depth and focused analyses. Starting with time-series gene expression data across *T* time points, DTPSP selects a subset *S* of critical time points that best capture the temporal dynamics of the system. The selected time points are then prioritized for more comprehensive profiling, such as single-cell sequencing or multi-omics studies (Fig. 1a). The input data, typically bulk RNA-seq due to its affordability and availability, undergoes preprocessing. This includes padding non-selected time points, adding time embeddings to incorporate temporal information, and integrating signals from neighboring genes. The processed data is then used by a deep learning model to predict gene expression across all time points. The model operates through a three-stage process: an initial pretraining phase where an autoencoder reconstructs the input data, a training phase where a regressor predicts gene expression for non-selected target time points, and a fine-tuning phase where the model is optimized to recover the full temporal trajectory. By evaluating all possible subsets of time points, DTPSP identifies the subset *S* that achieves the highest prediction accuracy, representing the most informative time points for downstream analyses (Fig. 1b).

**Fig. 1:**
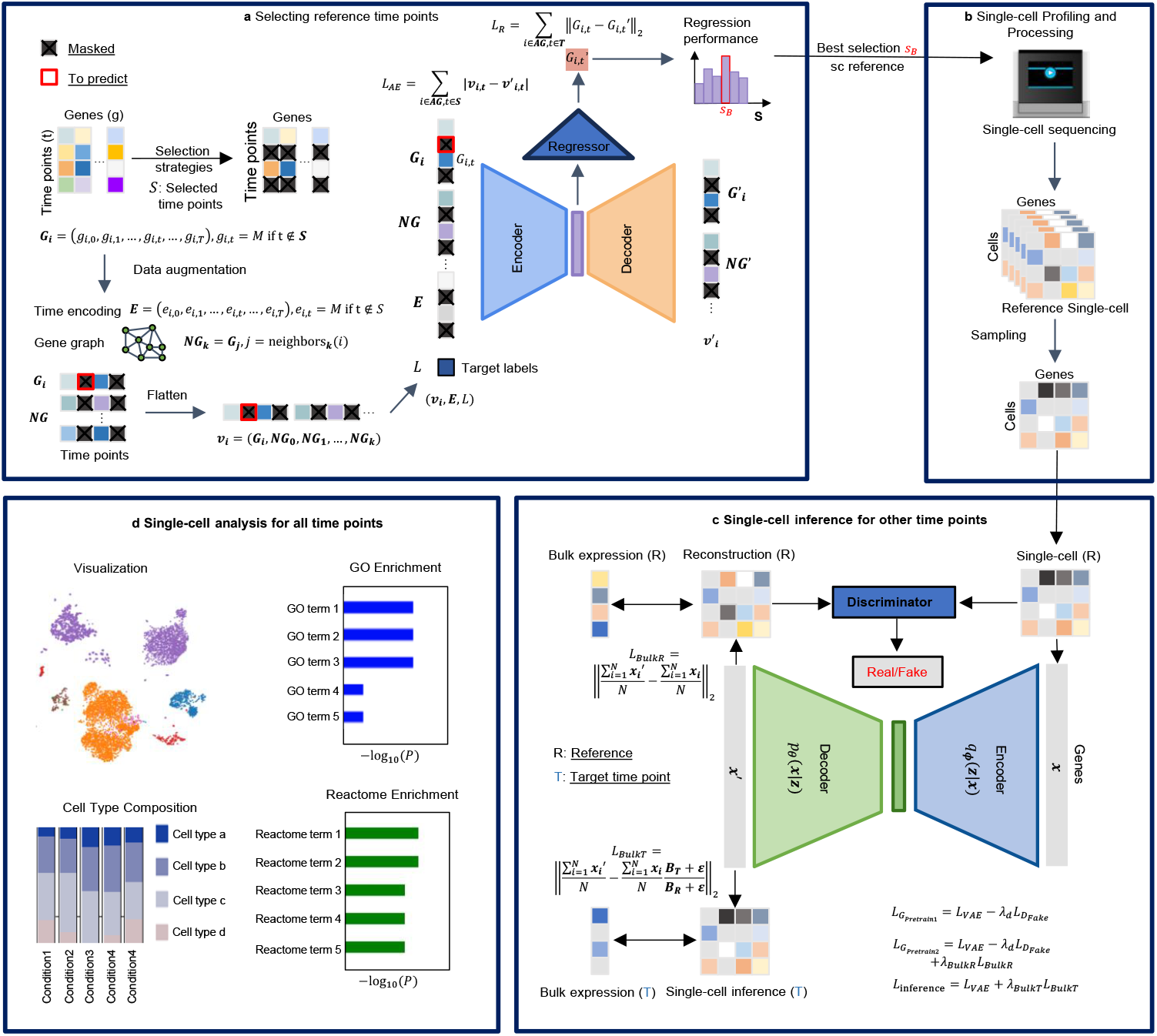
DTPSP method overview. a, Reference time point selection: A subset of time points, **S**, is selected from temporal bulk gene expression data spanning *T* time points and sent to a deep learning model to predict gene expression at non-selected time points. Gene expression data from **S** are augmented with neighboring genes (**NG**) with similar expression patterns, time embeddings (**E**), and a target label (*L*) that specifies the time point to be predicted. The deep regressor model undergoes pretraining with an autoencoder-based reconstruction task before being fine-tuned for regression to predict gene expression. The subset achieving the best prediction performance, **S**_**B**_, is identified as the most informative set of time points for subsequent analyses. **b, Single-cell profiling and processing:** The selected optimal time points, **S**_**B**_, are subjected to single-cell sequencing experiments to obtain real single-cell data. Standard single-cell processing pipelines are then applied to these data. **c, Single-cell inference for unselected time points (temporal single-cell semi-profiling):** For non-selected time points, a VAE-GAN model [25] is employed to infer single-cell data matrices. The VAE-GAN is first trained on a reconstruction task to learn the data distribution of the selected time points with real single-cell data. Subsequently, bulk gene expression data are incorporated into the model to transition the process from reconstruction to inference, generating single-cell data for the target time points. This results in a semi-profiled single-cell temporal trajectory where single-cell data are available for all time points. **d, Single-cell analysis across all time points:** The semi-profiled single-cell temporal trajectory enables comprehensive single-cell analyses, including visualization, deconvolution, enrichment analysis, and more.

In cases where single-cell resolution is required, DTPSP integrates a deep generative module to reconstruct single-cell profiles for non-sampled (unselected) time points. This module, based on a Variational Autoencoder-Generative Adversarial Network (VAE-GAN)[25] adapted from scSemiProfiler, utilizes the single-cell data from the selected time points as a reference and incorporates bulk differences between time points to infer single-cell profiles for non-sampled intervals. This process reconstructs the complete single-cell time-series dataset, enabling downstream analyses such as cell type composition estimation, biomarker discovery, and functional enrichment studies (Fig. 1c, 1d). DTPSP thus provides a flexible framework that generalizes across bulk and single-cell data. By leveraging affordable data sources for time point selection and prioritizing critical time points for in-depth profiling, DTPSP reduces experimental costs while enhancing biological insights through resource-efficient experimental designs and comprehensive temporal analyses.

### DTPSP Accurately Reconstructs Temporal Gene Expression Dynamics

To validate DTPSP’s ability to predict and select informative time points for capturing bulk-level temporal dynamics, we tested it on three publicly available datasets. Two datasets—(1) a lung alveoli dataset from Zepp et al. [26] and (2) the LungMap dataset [27]—consist exclusively of single-cell RNA-seq samples collected across seven time points each. From these single-cell datasets, pseudobulk profiles were generated by averaging normalized expression counts, creating bulk-like data suitable for DTPSP’s time-point prediction and selection tasks. Analyses were conducted on the top 6,000 highly variable genes identified from the pseudobulk data. The third dataset—(3) the iPSC-induced iMGL cell dataset from Ramaswami et al. [28]—includes both single-cell and bulk RNA-seq data measured at five time points, enabling direct validation with actual bulk data. For this dataset, we used the 6,013 highly variable genes identified in Wang et al.’s study [29].

By applying DTPSP to these datasets, we evaluated its ability to accurately predict missing time points and identify minimal subsets of time points that effectively capture temporal dynamics. DTPSP tested all possible subsets of time points to reconstruct temporal transcriptomic trajectories, with performance assessed by comparing reconstructed profiles to the ground truth for time points masked during inference. Figure 2a presents the average prediction performance—measured by mean absolute error (MAE) and *R*^2^ score—across the Lung Alveoli, LungMap, and iMGL datasets. For the iMGL dataset, the model was trained using selected time-point samples from one biological replicate, and prediction errors were computed against the actual gene expression from another biological replicate to validate robustness. The dashed line in Figure 2a denotes the theoretical optimal performance, derived by comparing two replicates. For the Lung Alveoli and LungMap datasets, technical replicates were generated by dividing the data into two halves. For the iMGL dataset, the two biological replicates provided direct comparisons. The star symbol marks the “elbow point”, representing the optimal number of selected time points that balance prediction accuracy with experimental cost.

**Fig. 2:**
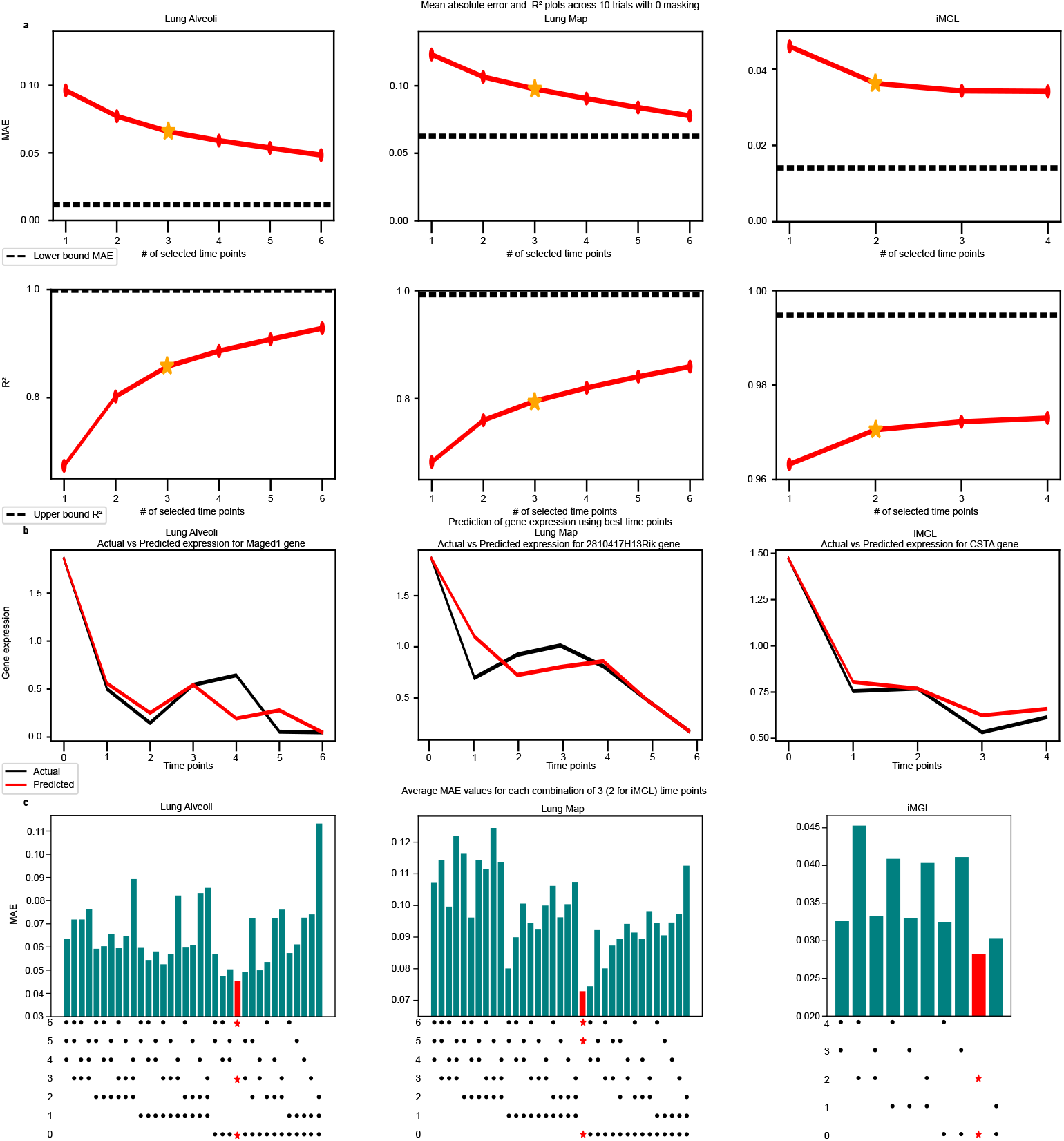
DTPSP selects the most informative reference time points based on accurate temporal trajectory prediction. **a, Prediction performance of DTPSP across three time-series bulk gene expression datasets**. The y-axis shows prediction performance (MAE or *R*^2^), and the x-axis indicates the number of reference time points selected. Dashed lines represent the theoretical best performance from technical or biological replicates. Yellow stars mark the “elbow point”, identifying the optimal number of time points that balance accuracy and cost. DTPSP achieves near-optimal accuracy as more time points are selected, consistently performing close to the theoretical maximum across datasets. **b, Reconstructed versus ground truth temporal trajectories for key biomarkers**. Red lines represent reconstructed trajectories predicted by DTPSP, and black lines indicate the ground truth. Temporal biomarkers are DEGs between the first two time points. Each column corresponds to a dataset, with selected time points at the elbow point (from **a**) used to predict gene expression at other time points. Reconstructed trajectories closely match the ground truth in trend and shape, demonstrating DTPSP’s accuracy. **c, Average MAE for combinations of selected time points**. For the Lung Alveoli and LungMap datasets, all combinations of three time points are evaluated, while two time points are tested for iMGL. Black points below each bar show selected time points for each combination. Red bars, marked with stars, indicate the combination with the lowest MAE. Optimal selections significantly outperform others, underscoring their suitability for accurate temporal trajectory prediction.

DTPSP demonstrated consistent and robust performance across all datasets. On the Lung Alveoli dataset, the model achieved impressive accuracy at the elbow point with three selected time points, reaching an average MAE of 0.066 and an *R*^2^ score of 0.86, closely approximating the theoretical performance. On the LungMap dataset, DTPSP achieved an MAE of 0.098 and an *R*^2^ score of 0.80 at the elbow point with three out of seven time points selected, with an MAE only 0.035 higher than the theoretical lower bound. These results highlight DTPSP’s robustness, even in pseudobulk scenarios. On the iMGL dataset, which includes actual bulk RNA-seq data, DTPSP delivered near-perfect performance, with an MAE of 0.035 and an *R*^2^ score of 0.97 at the elbow point. This superior performance compared to pseudobulk datasets is likely due to the higher signal quality of real bulk RNA-seq data, as single-cell data are inherently noisier, and their derived pseudobulk profiles contain less information. These findings demonstrate DTPSP’s capacity to accurately reconstruct temporal gene expression dynamics across diverse experimental settings.

Figure 2b illustrates reconstructed temporal trajectories for representative genes from each dataset. Genes were selected as top time-series markers based on their differential expression between the first two time points. From left to right, examples include *Maged1* (Lung Alveoli dataset), *2810417H13Rik* (LungMap dataset), and *CSTA* (iMGL dataset). In all cases, the predicted trajectories closely align with observed data, capturing both overall trends and precise expression levels. This demonstrates DTPSP’s capability to infer the dynamics of non-selected time points and reconstruct the temporal landscape of gene expression changes using a minimal subset of sampled time points.

The impact of selecting informative time points is evident in Figure 2c, where time points chosen by DTPSP (marked in red) yield consistently lower prediction errors. These selected time points are enriched for temporal information and serve as optimal candidates for in-depth profiling, such as single-cell sequencing or multi-omics studies, to capture cellular dynamics and heterogeneity. For the Lung Alveoli dataset, selecting three time points (time points 0, 3, and 6) resulted in the lowest MAE of 0.045 and an *R*^2^ score of 0.90. Similarly, for the LungMap dataset, the optimal three time points (time points 0, 5, and 6) achieved an MAE of 0.073 and an *R*^2^ score of 0.62, significantly outperforming all other selections. For the iMGL dataset, DTPSP achieved exceptional generalizability across biological replicates, with an MAE of 0.028 and an *R*^2^ score of 0.99 using only two time points (time points 0 and 2). These results emphasize DTPSP’s ability to consistently identify subsets of time points that effectively capture essential temporal dynamics, achieving high accuracy while minimizing experimental costs.

### DTPSP’s Bulk Expression Temporal Trajectory Prediction Supports Reliable Transcriptomic Analysis

To validate DTPSP’s reliability in predicting bulk temporal gene expression trajectories, we evaluated its performance on downstream transcriptomic analysis tasks using the Lung Alveoli dataset. By comparing results derived from predicted and real datasets, we assessed DTPSP’s ability to reconstruct gene expression at non-profiled time points and retain biological and technical reliability. In the predicted dataset, DTPSP reconstructed all genes at non-profiled time points using selected informative time points as a reference, creating a fully simulated dataset for analysis (Fig. 3).

**Fig. 3:**
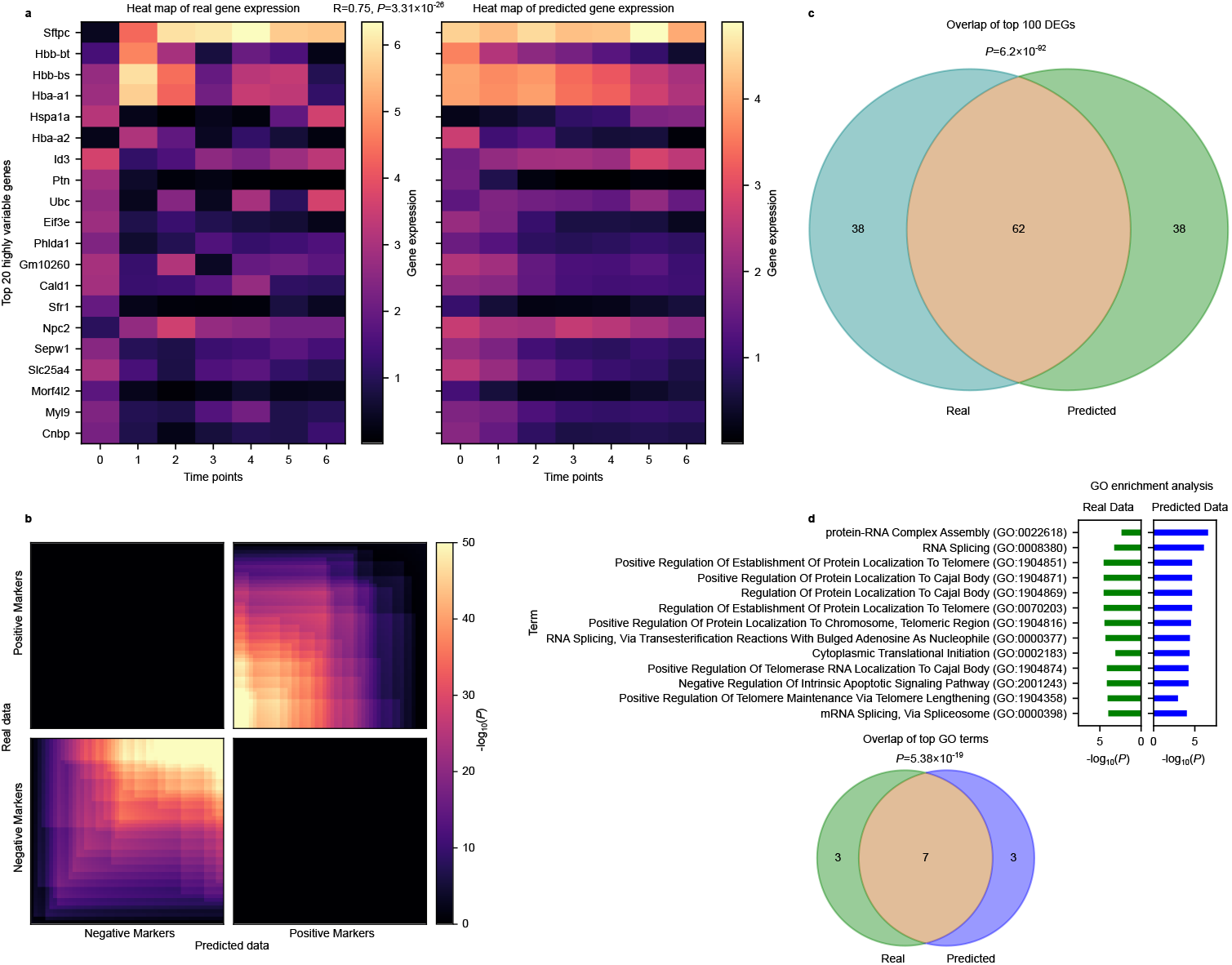
Biological validation of DTPSP predictions on the Lung Alveoli dataset. **a**, Heatmap comparison of real and predicted gene expressions. The top 20 differentially expressed genes (DEGs) with the lowest p-values, identified using the real dataset, are shown. In the predicted dataset (right heatmap), gene expressions at each time point are inferred using all other time points as input. Pearson correlation (*R* = 0.75) and p-value (*P* = 3.31 *×* 10^−26^) are calculated by flattening the heatmap matrices, demonstrating a strong correlation and visual similarity between real and predicted data. **b**, Rank-Rank Hypergeometric Overlap (RRHO) plot illustrating the overlap of positive and negative marker genes between the real and predicted datasets. Bright regions in the first and third quadrants indicate significant concordance of markers, supported by hypergeometric p-values in these regions. **c**, Venn diagram showing the overlap of the top 100 DEGs identified from the real and predicted datasets. A significant overlap of 62 genes is observed, with a hypergeometric p-value of 6.2 *×* 10^−92^, validating the reliability of DTPSP in recovering biologically relevant DEGs. **d**, GO enrichment analysis comparing the real and predicted datasets. The union of the top 10 Gene Ontology (GO) terms enriched in each dataset is shown. A Venn diagram highlights a significant overlap of 7 terms, with a hypergeometric p-value of 5.38 *×* 10^−19^, confirming the biological consistency of DTPSP’s predictions for pathway enrichment analysis.

We first compared differential gene expression (DEG) analysis results between the predicted and real datasets. DEGs were identified between time points 0 and 1 in the real dataset using ground truth values at both time points, while the predicted dataset used real values at time point 0 and reconstructed values at time point 1. Heatmaps of the top 20 DEGs (Fig. 3a) reveal highly similar expression patterns, with the top four DEGs showing consistently elevated expression levels in both datasets. The similarity is quantified by a Pearson correlation coefficient of 0.75 (p-value = 3.31 *×* 10^−26^). Additionally, comparing the top 100 DEGs from the two datasets (Fig. 3b) shows a significant overlap of 62 genes (hypergeometric test p-value = 6.2 *×* 10^−92^), demonstrating DTPSP’s ability to recover biologically relevant DEGs from reconstructed time points. To assess biomarker discovery, we compared marker genes identified in the real and predicted datasets using a Rank-Rank Hypergeometric Overlap (RRHO) [30] plot (Fig. 3c). The top 50 positive and top 50 negative markers showed substantial overlap, with bright regions in the first and third quadrants indicating significant concordance between datasets (supported by hypergeometric p-values). The dark regions in the second and fourth quadrants confirmed the expected lack of overlap between positive and negative markers. These results highlight DTPSP’s ability to preserve the rankings and biological relevance of marker genes.

We further evaluated biological insights from pathway enrichment analysis by comparing Gene Ontology (GO) [31, 32] terms enriched in biomarkers from the real and predicted datasets. Figure 3d shows a Venn diagram of the top 10 enriched terms, with 7 terms overlapping between the two datasets. This substantial overlap underscores DTPSP’s capability to capture critical biological pathways, validating its reliability for downstream transcriptomic analyses.

In summary, the results in Fig. 3 demonstrate that downstream transcriptomic analysis based on the predicted time-series trajectory generates highly similar results with the real counterpart. Similar analysis performance on the LungMap and iMGL datasets with real bulk data are presented in Supplementary Fig. S1 and S2, respectively. Overall, these results demonstrate that DTPSP not only faithfully reconstructs temporal gene expression trajectories at non-profiled time points but also preserves biological relevance for transcriptomic analyses. By supporting reliable differential gene expression identification, biomarker discovery, and pathway enrichment analysis, DTPSP facilitates robust and biologically meaningful insights in high-throughput time-series studies.

### DTPSP Advances Single-Cell Resolution Time-Series Studies

DTPSP not only optimally selects time points for time-series studies but also generates single-cell resolution data for unselected time points using a deep generative model (VAE-GAN [25]). This process, referred to as “single-cell inference”, reconstructs the entire time-series single-cell dataset (“time-series semi-profiling”), providing a cost-effective solution for single-cell time-series analysis. By aligning inferred data closely with real single-cell measurements, DTPSP ensures data reliability while reducing the need for exhaustive and expensive profiling across all time points.

To demonstrate DTPSP’s ability to generate single-cell resolution time-series data, we applied the scSemiProfiler framework to the LungMap dataset. Using optimal reference time points selected by DTPSP, scSemiProfiler reconstructed single-cell profiles for unprofiled time points by integrating bulk data and reference single-cell profiles. This approach retained high-resolution data at reduced experimental costs. Comparisons with a baseline clustering-based reference selection approach showed that DTPSP-selected time points served as a more informative scaffold for downstream single-cell inference, effectively capturing temporal dynamics and cellular heterogeneity.

The inferred single-cell data closely mirrored the real profiles, as shown in UMAP projections (Fig. 4a–c). Semi-profiled cells (Fig. 4b) exhibited clustering patterns nearly identical to the real cells (Fig. 4a), preserving the structure and spacing of cell populations. Notably, no artificial clusters were introduced, and the arrangement of inferred cells faithfully recapitulated transitions and relative positions observed in the real data (Fig. 4d). This demonstrates DTPSP’s ability to accurately reconstruct cellular complexity across time points, ensuring reliable representation of both global trends and finer transcriptional nuances.

**Fig. 4:**
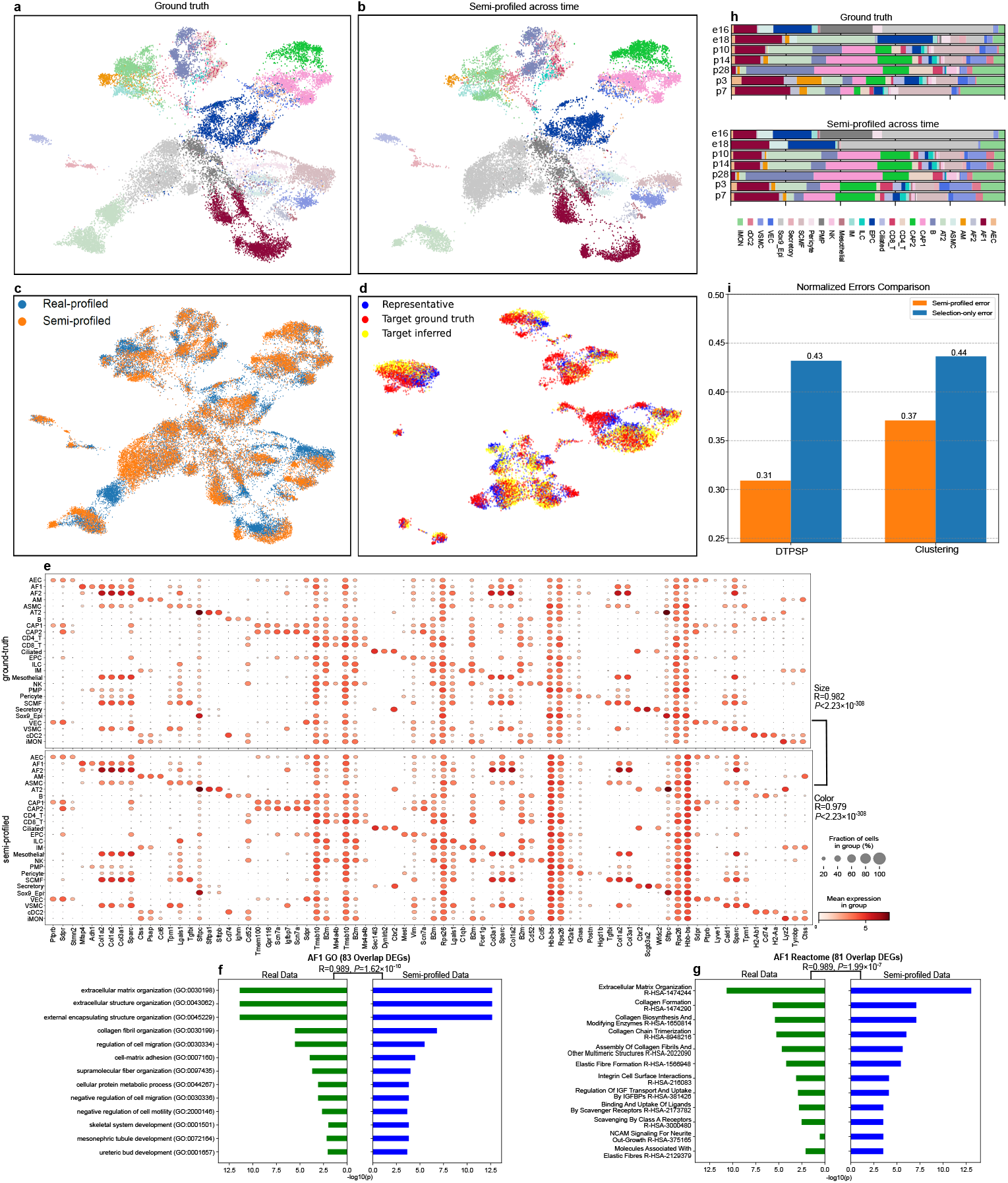
Comparison of semi-profiled and real-profiled LungMap datasets. **a**, UMAP visualization of the real-profiled dataset, with cell types colored as in (**h**). **b**, UMAP visualization of the semi-profiled dataset, showing clustering patterns similar to the real-profiled data. **c**, Integrated UMAP visualization of semi-profiled and real-profiled datasets, where distinct colors represent the data source. Overlapping regions indicate close alignment between the two datasets. **d**, UMAP visualization of cells at time point P10, demonstrating the fidelity of the semi-profiled data to real measurements. **e**, Dot plots comparing cell type-specific expression patterns in real-profiled (top) and semi-profiled (bottom) datasets. Dot size represents the fraction of cells expressing a gene, and color indicates average expression intensity. P-values from Pearson correlation tests highlight strong concordance between the datasets. **f**, GO term enrichment analysis of AF1 signature genes in both datasets. Bars represent the top 10 enriched terms, with lengths reflecting significance (Pearson correlation of p-values). **g**, Reactome pathway enrichment analysis of AF1 signature genes in both datasets. Bar lengths and significance levels are presented similarly to (**f**), showing a high degree of similarity. **h**, Stacked bar plot comparing cell type proportions across all time points. The upper portion represents real-profiled data, and the lower portion represents semi-profiled data, demonstrating consistent cell type composition. **i**, Bar plot showing that DTPSP-selected time points result in lower semi-profiling and selection-only errors compared to clustering-based selection.

At the individual sample level, inferred profiles closely aligned with real single-cell data, validating the effectiveness of DTPSP-selected reference points. Inferred cells not only overlapped with reference samples in transcriptional space but also migrated toward target sample profiles, capturing unique temporal and transcriptional cues. This local fidelity highlights the ability of DTPSP-selected time points to provide a biologically relevant scaffold for inference, enabling accurate reconstruction of single-cell resolution data for samples previously represented only by bulk measurements.

Biological fidelity was further validated through cell type-specific analyses. Dot plots of gene expression patterns (Fig. 4e) revealed a high correlation (*R* = 0.982, *p <* 2.23 *×* 10^−308^) between real and inferred profiles. Marker gene expression patterns for all cell types were preserved. These results confirm the utility of DTPSP in generating biologically meaningful single-cell data for time-series studies.

Pathway enrichment analyses provided additional evidence of biological consistency. GO and Reactome analyses of the top 100 cell type-specific signature genes revealed significant overlap between real and inferred data (Fig. 4f, g). Terms such as ‘extracellular matrix organization’ and ‘regulation of cell migration’ aligned with Reactome pathways like collagen formation and integrin signaling. The Pearson correlation coefficient for Reactome term significance levels was 0.989 (*p* = 1.62 *×* 10^−10^), further validating the inferred data’s ability to capture critical biological pathways and processes.

In Fig. 4h, stacked bar plots illustrate how cell type proportions shift over time in both the real-profiled and semi-profiled time-series datasets. The close correspondence between these two distributions indicates that DTPSP effectively captures the evolving cellular composition of the LungMap dataset, accurately reflecting heterogeneity and dynamic changes in cell populations across successive time points. This justifies DTPSP’s reliability for cell-type-level analysis.

To quantify the advantage of DTPSP over clustering-based reference selection, we computed error metrics assessing how closely inferred profiles aligned with real data. DTPSP-selected representatives yielded lower semi-profiled errors (normalized Euclidean distance) than clustering-based selections (Fig. 4i). Additionally, DTPSP consistently achieved a larger improvement in accuracy from semi-profiling compared to substituting target samples with representatives directly. These results demonstrate that DTPSP provides a superior scaffold for single-cell inference, enabling accurate reconstruction of cellular dynamics and advancing single-cell resolution time-series studies.

### DTPSP maintains high fidelity in highly heterogeneous time-series dataset

We further evaluated the DTPSP approach on the more challenging Lung Alveoli dataset (Fig. 5). Unlike the LungMap dataset, the Lung Alveoli samples exhibit significantly greater heterogeneity across time points. This reflects the dynamic transcriptional states and structural remodeling in the alveolar region during lung development, driven by the interplay of mesenchymal, epithelial, and endothelial cells [33]. This greater heterogeneity poses a more complex single-cell inference task due to the distinct cellular distributions observed across developmental stages. Despite these increased challenges, the semi-profiled data closely matched the real single-cell data. UMAP projections (Fig. 5a–c) demonstrate that the inferred cells accurately recapitulated the underlying cellular landscape, capturing spatial patterns and cluster boundaries that mirror those of the ground truth data. Examining individual samples at high resolution (Fig. 5d), the inferred target cells positioned themselves accurately within the target sample’s local transcriptional space rather than merely clustering around the representative samples. This indicates that even in the presence of high heterogeneity, the DTPSP-identified representatives provided a robust scaffold for semi-profiling, enabling the deep generative learning component to generate high-fidelity, sample-specific single-cell profiles for cells not directly measured.

**Fig. 5:**
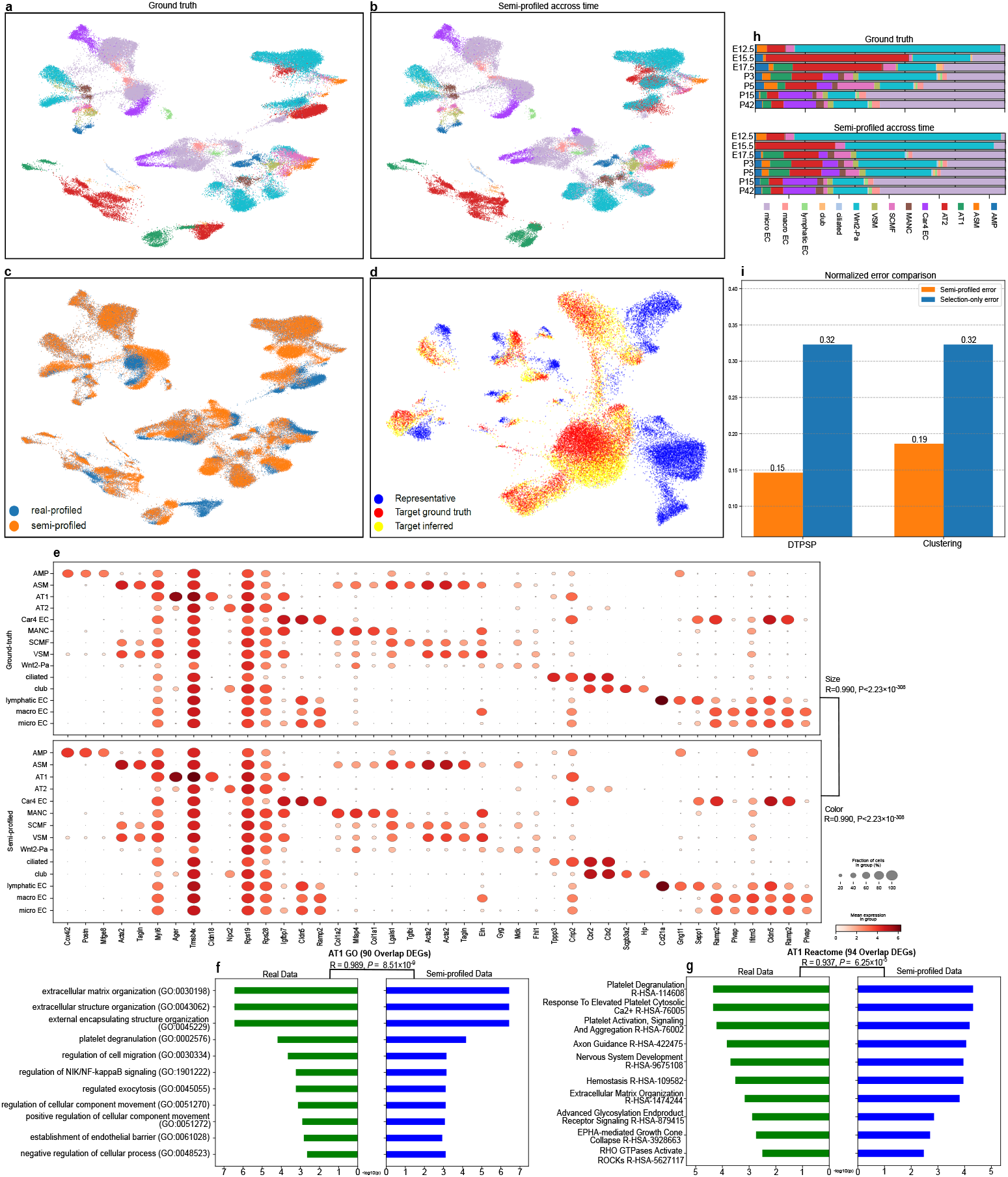
Overall comparisons of the semi-profiled and real-profiled Alveolus dataset. **a** UMAP visualization of the real-profiled data. Colors correspond to cell types and are consistent with (**h**). **b** UMAP visualization of the semi-profiled data. **c** Integration of both time-series semi-profiled and real-profiled datasets in a UMAP visualization, with distinct colors indicating the data source of each cell. Overlapping regions highlight areas where the semi-profiled data closely align with the real-profiled data. **d** UMAP visualization of cells at time point P15, highlighting the representative sample (blue), inferred target sample cells (yellow), and real single-cell profiles of the target sample (red). **e** Dot plots showing expression proportion and intensity of cell type signature genes. The top half displays the real-profiled dataset, while the bottom half shows the semi-profiled version. P-values are from two-sided Pearson correlation tests. **f** GO term enrichment based on AT1 signature genes from both datasets. The union of the top 10 enriched terms is highlighted, with bar lengths reflecting Pearson correlation coefficients from corresponding p-values. **g** Reactome enrichment based on AT1 signature genes from both datasets. The top 10 enriched terms are shown with bar lengths and p-values as described above. **h** Stacked bar plot illustrating the proportions of cell types across all timepoints. The upper portion represents real-profiled data, while the lower portion depicts semi-profiled data. **i** Bar plot comparing normalized semi-profiled and naive errors across methods.

Cell-type-specific gene expression patterns and enrichment analyses in the Lung Alveoli dataset further support the robustness of our approach (Fig. 5e–g). The dot plots(Fig. 5e) comparing real and semi-profiled data indicate a highly consistent representation of cell type marker genes, both in terms of the fraction of cells expressing a given gene (dot size) and the average expression level (dot color). Pearson correlations exceeding 0.99 (*P <* 2.23 *×* 10^−308^) attest to the remarkable fidelity of our inferred profiles. Focusing on alveolar type 1 (AT1) cells as an example, the sets of top differentially expressed genes (DEGs) identified from real and semi-profiled data showed substantial concordance (Fig. 5f, g). Specifically, 90 out of the top 100 AT1 DEGs identified in real data also appeared in the semi-profiled dataset’s top 100 DEGs, and 94 overlapping DEGs were found when considering Reactome pathways. Functional enrichment of these DEGs revealed highly similar key biological processes and pathways. For instance, the Pearson correlation coefficient between the - log_10_(*p*) values of the top 10 significantly enriched GO terms from the real and semi-profiled data was 0.989 (*P* = 8.51 *×* 10^−9^), while the correlation for the top 10 Reactome terms was 0.937 (*P* = 6.25 *×* 10^−5^). This quantitative alignment confirms that not only are individual gene-level patterns accurately reconstructed, but also that the broader functional landscape is faithfully recovered.

In addition, Fig. 5h shows stacked bar plots depicting cell type proportions over time in both the real-profiled and semi-profiled Lung Alveoli datasets. The strong similarity between these distributions indicates that DTPSP accurately captures the evolving cellular composition, faithfully reflecting heterogeneity and dynamic shifts in cell populations across successive time points. This demonstrates DTPSP’s reliability for cell-type-level analyses.

Remarkably, for the Lung Alveoli dataset, the representative sets chosen by DTPSP and the clustering-based method differed by only one time point: DTPSP selected E12.5, whereas clustering selected E15.5, with both methods also choosing P3 and P42. Since our selection-only error computation is symmetric—effectively measuring how well each chosen representative sample alone approximates any given target sample—replacing E15.5 with E12.5 does not change the selection-only error. However, this subtle shift dramatically reduced the semi-profiled error (Fig. 5i). This improvement underscores the power of DTPSP: by pinpointing a slightly earlier developmental stage (E12.5) as a reference, DTPSP provides a more suitable temporal anchor for semi-profiling. As a result, the deep generative learning component can more accurately capture early transcriptional states and better reconstruct the trajectories of target samples. Thus, even when representative sets appear similar, DTPSP’s data-driven optimization identifies the optimal choice of time points, yielding a significant improvement in the fidelity of inferred single-cell profiles.

Overall, the performance of DTPSP on the Lung Alveoli dataset demonstrates its efficacy in handling highly heterogeneous and complex time-series data. The ability to maintain lower normalized errors compared to the LungMap dataset, despite the increased variability, highlights DTPSP’s robustness and adaptability in diverse biological contexts. This ensures that even in scenarios with significant sample dissimilarity, DTPSP-selected representatives provide a reliable foundation for accurate single-cell inference, facilitating comprehensive biological insights across varying developmental and pathological states.

### DTPSP Demonstrates Reliability with Real Bulk RNA-seq Data: Application to the iMGL Dataset

In the previous sections, we demonstrated the informativeness of DTPSP’s time point selection and its ability to achieve highly accurate inference of single-cell time-series trajectories. However, these results were based on datasets with pseudobulk data, generated by averaging single-cell data. While informative, this does not fully justify DTPSP’s performance in real-world scenarios where real bulk sequencing data is used. This limitation arises due to systematic differences between single-cell and real bulk data. For example, single-cell data tend to be noisier and exhibit higher variability than bulk data, leading to discrepancies between real bulk data and pseudobulk profiles derived from single-cell data[16, 34]. To address this and evaluate DTPSP’s capability in selecting informative time points and guiding single-cell trajectory reconstruction in a real-world setting, we applied our method to the iMGL dataset, which includes both real bulk and real single-cell ground truth. Unlike the LungMap and Lung Alveoli datasets, which are based on pseudo-bulk data derived from single-cell experiments, the iMGL dataset comprises genuine bulk RNA-seq samples. This allowed us to test whether DTPSP could successfully select representative time points and generate high-fidelity single-cell profiles directly from authentic bulk data, thereby demonstrating its utility in more practical and widely applicable settings. Figures 6a–c illustrate that the semi-profiled cells closely align with the real-profiled cells at a global level. The UMAP projections of the semi-profiled and real-profiled data strongly resemble each other, exhibiting similar cluster shapes and distributions. When overlaid (Fig. 6c), the inferred cells occupy the same regions of the transcriptional landscape as the real cells, indicating that the temporal dynamics and cellular heterogeneity are well-captured. This close similarity is further exemplified at the individual sample level (Fig. 6d), where semi-profiled target cells based on DTPSP-chosen references more closely approximate the real single-cell profiles than the representative sample alone. Despite relying on actual bulk data instead of pseudo-bulk averages, the inference still accurately reconstructs sample-specific transcriptional states, underscoring the robustness of the approach.

**Fig. 6:**
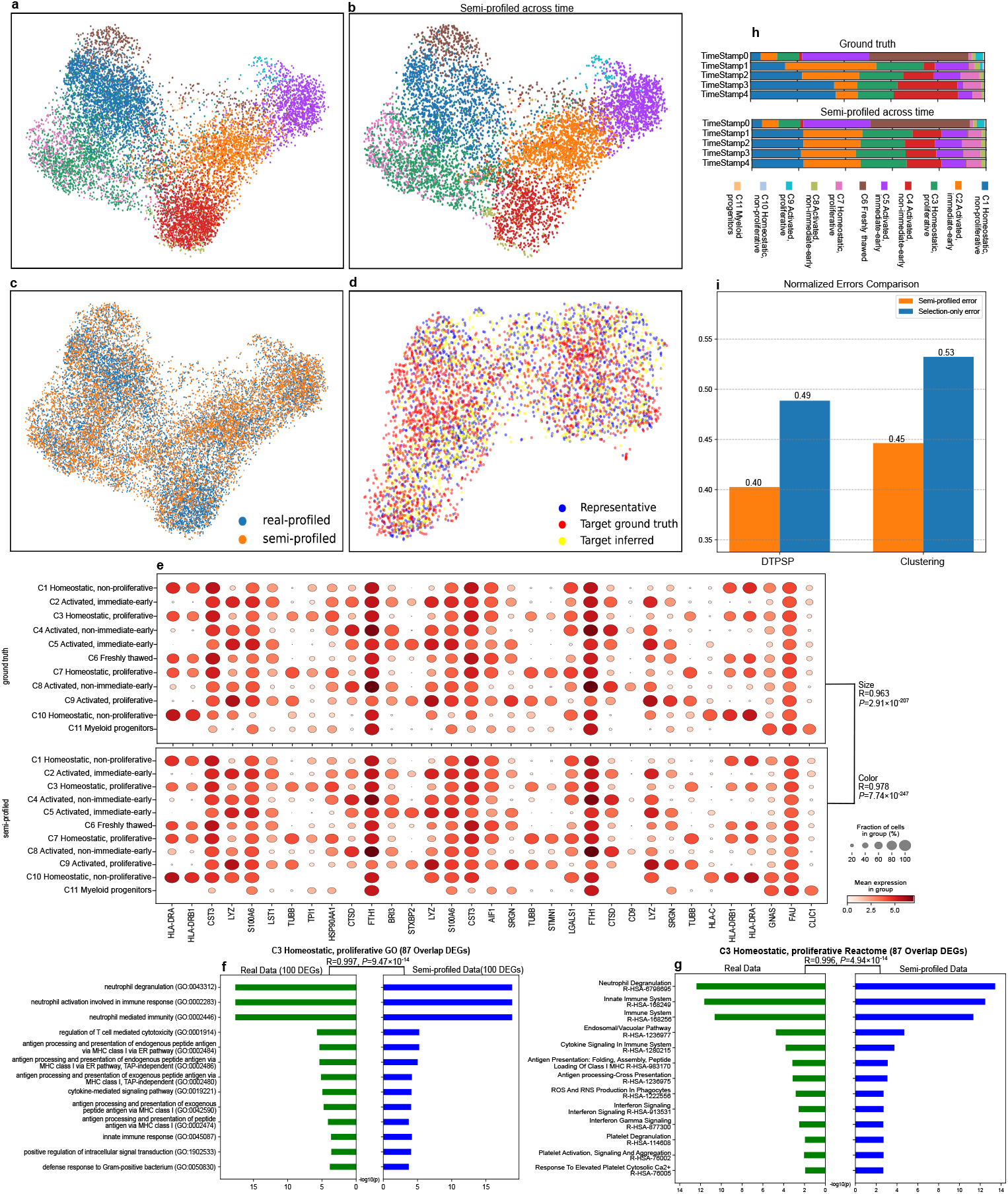
Overall comparisons of the semi-profiled and real-profiled iMGL dataset. **a** UMAP visualization of the real-profiled data. Colors correspond to cell types and are consistent with (**h**). **b** UMAP visualization of the semi-profiled data. **c** Integration of both time-series semi-profiled and real-profiled datasets in a UMAP visualization, with distinct colors indicating the data source of each cell. Overlapping regions highlight areas where the semi-profiled data closely align with the real-profiled data. **d** UMAP visualization of cells at time point Day1. **e** Dot plots showing expression proportion and intensity of cell type signature genes. The top half displays the real-profiled dataset, while the bottom half shows the semi-profiled version. P-values are from two-sided Pearson correlation tests. **f** GO term enrichment based on C3 Homeostatic, proliferative Reactome signature genes from both datasets. The union of the top 10 enriched terms is highlighted, with bar lengths reflecting Pearson correlation coefficients from corresponding p-values. **g** Reactome enrichment based on C3 Homeostatic, proliferative Reactome signature genes from both datasets. The top 10 enriched terms are shown with bar lengths and p-values as described above. **h** Stacked bar plot illustrating the proportions of cell types across all timepoints. The upper portion represents real-profiled data, while the lower portion depicts semi-profiled data. **i** Bar plot comparing normalized semi-profiled and naive errors across methods.

Examining cell-type-specific markers and proportions reveals high fidelity between real and semi-profiled data. The dot plots in Fig. 6e show a strong similarity in both the fraction of cells expressing key genes (indicated by dot size) and their average expression levels (indicated by dot color). The Pearson correlation for the fraction of cells expressing these genes is *R* = 0.963 with *P* = 2.91 *×* 10^−207^, while the correlation for mean expression levels is even higher at *R* = 0.978 with *P* = 7.74 *×* 10^−247^. This robust agreement confirms that the biological identity and functional characteristics of the cell populations are well-preserved. Importantly, these results demonstrate that rare or transient cell states of interest are accurately inferred and not lost during the semi-profiling process.

In terms of functional enrichment, gene sets identified from the top differentially expressed genes (DEGs) demonstrate a strong concordance between the real and semi-profiled datasets. As shown in Fig. 6f, GO terms related to key biological processes, such as ‘neutrophil degranulation’, ‘neutrophil activation involved in immune response,’ and ‘regulation of T cell-mediated cytotoxicity,’ exhibit highly similar enrichment scores, with a Pearson correlation of *R* = 0.997 and *P* = 9.47 *×* 10^−14^. Similarly, Fig. 6g highlights the overlap in Reactome pathways, including ‘Neutrophil Degranulation’, ‘Innate Immune System,’ and ‘Cytokine Signaling in Immune System’, with a Pearson correlation of *R* = 0.996 and *P* = 4.94 *×* 10^−14^. These consistent enrichment scores confirm that both the cellular identity and the broader functional contexts are faithfully recovered when using semi-profiled data derived from real bulk RNA-seq data.

In Fig. 6h, we present the cell type deconvolution results for the iMGL dataset, which includes real bulk RNA-seq data. This task is particularly challenging given that the cells are all iPSC-induced iMGL subtypes, resulting in a high degree of homogeneity and making it more difficult to distinguish subtle changes in cell composition. Nevertheless, informed by DTPSP’s optimal time point selection, we accurately reconstruct the single-cell time-series data, yielding stacked bar plots of cell type proportions closely matching those of the ground truth. These results underscore DTPSP’s ability to reliably capture evolving cellular landscapes, even in highly homogeneous populations, and confirm its utility for cell-type-level analyses in realistic experimental settings.

Finally, as shown in Fig. 6i, using DTPSP-derived representatives yields substantially lower semi-profiled errors than naive clustering-based choices. Specifically, the selection-only error for DTPSP (0.49) is smaller than for the clustering method (0.53). The difference in semi-profiling error is even more significant, with DTPSP achieving an error of 0.40 compared to 0.45 for the clustering approach. This improvement in inference accuracy underscores the generalizability of DTPSP’s reference selection strategy, even in the setting of true bulk data. By focusing on time points that comprehensively capture the underlying temporal dynamics selected by DTPSP’s regressor component, DTPSP ensures that its deep generative learning component can reconstruct single-cell states with high fidelity from a cost-effective sampling strategy.

## Discussion

In this study, we introduced Deep Time Point Selector and Profiler (DTPSP), a deep learning-based framework designed to optimize and enhance single-cell time-series transcriptomics studies. DTPSP addresses the critical challenges of cost and feasibility in profiling single-cell data across multiple time points by first selecting the most informative time points using a supervised learning model and then reconstructing complete single-cell trajectories through a deep generative model. By combining bulk or pseudobulk data with deep learning, DTPSP provides high-resolution temporal data, enabling downstream analyses with high fidelity. Validation on three publicly available datasets demonstrated the robustness and reliability of DTPSP in identifying critical time points and generating reconstructed data that closely align with real-profiled single-cell data.

DTPSP offers several significant contributions to the field. First, it introduces a novel deep learning framework specifically designed to identify the most informative time points in time-series studies. Recognizing that not all time points are equally important, DTPSP optimizes resource allocation by focusing on key time points that capture critical temporal dynamics. This capability enables researchers to conduct more in-depth analyses, such as single-cell profiling or multi-omics studies, at these prioritized intervals, enhancing the resolution and depth of biological insight. Second, DTPSP provides a systematic and cost-efficient approach for experimental design. By leveraging accessible bulk or pseudobulk data for time-point selection, it minimizes the need for comprehensive profiling across all time points, significantly reducing experimental costs while preserving the ability to capture essential biological processes. This approach ensures that researchers can balance experimental feasibility with scientific rigor. Third, DTPSP extends beyond traditional methods of time-series analysis by integrating gene-gene relationships into its predictive framework. Through k-nearest neighbor (KNN) integration, DTPSP models complex gene interdependencies, improving its ability to predict temporal trajectories accurately. This feature makes the framework particularly suited for datasets with intricate temporal dynamics. Finally, DTPSP employs a deep generative model to reconstruct single-cell gene expression data at non-profiled time points. This capability bridges bulk and single-cell modalities, allowing researchers to gain high-resolution insights without the need for exhaustive single-cell sequencing. Together, these contributions position DTPSP as a versatile and transformative tool for optimizing time-series studies and uncovering dynamic biological processes. Despite its strengths, DTPSP has some limitations and presents opportunities for future development. Currently, the framework has been validated exclusively on RNA-seq data. Its application to multi-omics datasets, such as proteomics or epigenomics, remains unexplored. Extending DTPSP to multi-omics studies would enable more comprehensive analyses by integrating different molecular layers, providing deeper insights into complex biological systems. Additionally, while DTPSP has shown strong performance in temporal studies, its adaptability to diverse biological systems, such as disease progression or therapeutic response, requires further testing. Incorporating domain-specific knowledge or refining masking strategies to better account for temporal dependencies could enhance the model’s robustness and generalizability.

DTPSP provides a practical solution for time-series studies by identifying the most informative time points for focused and detailed analysis. By prioritizing key intervals, it facilitates efficient resource allocation for in-depth profiling, such as multi-omics or single-cell studies, while minimizing redundancy. This approach supports the study of dynamic systems, including development, disease progression, and therapeutic responses, by ensuring critical temporal dynamics are effectively captured. DTPSP offers a scalable and versatile framework that contributes to more efficient experimental designs in systems biology.

## Methods

### Public datasets used for validating DTPSP

The iMGL dataset comprises single-cell RNA sequencing and bulk RNA sequencing data from human-induced pluripotent stem cell (iPSC)-derived microglia-like cells. It includes 25 samples collected across different culture durations. For model training and testing, we selected 10 untreated samples from two replicates spanning time points from day 0 (D0) to day 4 (D4). One replicate was used to train the deep regressor model, while the predicted gene expression values were validated using the other replicate. This dataset was utilized to validate the capability of the proposed methods to generate single-cell data from real bulk RNA-seq profiles. The LungMAP dataset comprises single-cell RNA sequencing (scRNA-seq) data derived from mouse lungs at seven developmental time points: E16, E18, P3, P7, P10, P14, and P28. This dataset captures the dynamic changes in lung cell populations, including epithelial, endothelial, and mesenchymal cells, across critical stages of embryonic and postnatal development. It serves as a valuable resource for exploring the cellular and molecular transitions underlying lung maturation during these key developmental periods. The Lung Alveoli dataset provides scRNA-seq data from mouse lungs collected at seven developmental time points: E12.5, E15.5, E17.5, P3, P5, P15, and P42. This dataset focuses on the process of alveolar development, known as alveologenesis, and includes data on various cell types involved in this process, such as alveolar type 1 (AT1) and alveolar type 2 (AT2) cells, as well as mesenchymal lineages like secondary crest myofibroblasts (SCMFs). Spanning both embryonic and early postnatal stages, the dataset offers a detailed view of the dynamic cellular and molecular processes that drive the emergence and maturation of the lung alveolus.

### Data preprocessing

The standard SCANPY[35] preprocessing pipeline is applied to single-cell datasets: filtering low quality cells (less than 100 genes expressed) and low quality genes (expressed in fewer than 3 cells) for data from each time point. Only genes appearing in all time points are used for subsequent analysis. The data’s library size is normalized to 10,000 per cell. Subsequently, the data is thresholded: extremely low counts (less than 10 by default) are converted to zero because they are likely to be background noise according to previous studies [36, 37]. After the thresholding, library size is normed back to 10,000. To obtain the pseudobulk data, the normalized count of each sample is averaged across the cells. The pseduobulk data is then log1p transformed. SCANPY’s highly variable genes tool is applied to the log1p transformed pseudobulk data to select the top 6,000 highly variable genes (HVGs) from the bulk time-series dataset. In the case of iMGL dataset with real bulk data, the normed, log1p transformed real bulk data is used for selecting 6,000 highly variable genes. After HVGs are selected, only HVGs of the bulk and single-cell data are used. For the single-cell inference of iMGL dataset, we used the same 6,013 HVGs from scSemiProfiler’s study [29].

For training the deep regressor model, 80% of the genes are randomly selected for training and the rest are for testing. We prepared the input features by concatenating the data **G**_**i**_, a gene expression vector across time for gene *i* (*g*_*i*_), with neighbor genes **NG**_**i**_, time embeddings **E**, and the target label *L*. Let **v**_**i**_ denote the vector of (**G**_**i**_, **NG**_**0**_, **NG**_**1**_, …, **NG**_**k**_), 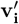 denote the reconstruction of **v**_**i**_ using the autoencoder, 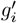 denote the predicted gene expression vector of gene *i* across time, **S** denote the set of selected (not masked) time points, *T* denote the largest time points value starting from 0, **E** is the time embeddings (*e*_*i*,0_,*e*_*i*,1_,…,*e*_*i,T*_) which are the integers (0,1,…,*T*), and *M* denote the masking value for not selected time points (0 by default). Neighbor genes are genes with similar expressions across all time points using Python package Scikit-learn’s[38] K-Nearest Neighbors function `sklearn.neighbors.NearestNeighbors`, with K being a hyperparameter set to 15 for Lung Alveoli, LungMap, and 25 for iMGL. Target label *L* is an integer specifying the target time point to be predicted: *L* ∈ **T, T** = *{*0, 1, 2, …, *T }*

### The architecture of the deep regressor for bulk temporal trajectory reconstruction

DTPSP is a hybrid model made up of two components, a deep regressor model for time-series trajectory reconstruction and time point selection, and a deep generative learning model for single-cell inference task adapted from scSemiProfiler. The deep regressor model consists of an autoencoder and a regressor subnetwork built using PyTorch[39]. The autoencoder consists of 2 Multi-layer Perceptrons (MLP), an encoder and a decoder. This input goes through a neural network with 4 layers: input layer, 2 hidden layers of 128 nodes each with one ReLU[40] activation function between them, and an output layer with the number of nodes equal to 64 for each dataset. The decoder has the exact same architecture, flipped, with a ReLU activation function after the output, to ensure the model does not predict negative values when reconstructing the gene expressions from the latent space, since there are no negative values in the data. The regressor subnetwork’s architecture is also an MLP with 4 layers, with 64, 64, 32, and 1 nodes, respectively, and ReLU between each layer and one at the end, and a Dropout layer, set to 0.2, before the last layer to avoid overfitting. The extra ReLU at the end ensures the model does not predict negative gene expression values. The input to the regressor subnet is directly from the latent space representation provided by the encoder.

### Deep regressor model training

The deep regressor’s training for predicting other time points’ gene expression value comprises three stages: (1) a pretraining stage using the autoencoder to reconstruct the input time points’ gene expression data, (2) a training stage affecting the encoder and the regressor subnetwork for predicting gene expression (3) a fine-tuning stage training both parts jointly. All stages are trained until convergence. The overall training objective can be represented by minimizing the loss function 1 below.

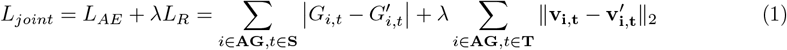

During the pretrain stage, the autoencoder aims to reconstruct the gene expression data of the selected time points and minimize the reconstruction loss *L*_*AE*_ in Eq. 1. This helps the model learn a encoder capable of extracting essential information from the input, getting the model prepared for the subsequent training stages. In *L*_*AE*_, *t* is time point in **T** = {0, 1, 2, …, *T* }, the total time points in the time-series dataset. **AG** is the set of all analyzed genes. **v**_**i**_ = (**G**_**i**_, **NG**_**0**_, …, **NG**_**k**_) is the gene expression feature corresponding to gene *i*. **NG**s are *k* neighbor genes for augmenting the features. **G** is the vector including all gene expressions across all time points and **NG** is the values for neighbor genes. In **v**_**i**_ only the gene expression at the selected time points in **S** have actual values while non-selected time points’ values are masked. **v**_**i,t**_ is the part of the input feature that corresponds to gene expressions at time point *t*. In short, the loss minimizes the difference between the ground truth and reconstructed gene expression of gene expressions at the selected time points. Adam optimizer[41] is used for the autoencoder and a learning rate of 1 *×* 10^−3^ and weight decay[42] of 1 *×* 10^−4^ to avoid overfitting.

After the pretraining stage, the model trains the regressor subnetwork, focusing on accurate prediction of the non-selected time points by minimizing the regression loss *L*_*R*_ in Eq. 1. *g*_*i,t*_ and 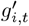 are ground truth and predicted gene expression of gene *i* time point *t* respectively. The predicted value is output by the regressor subnetwork taking the latent space representation generated by the encoder as input. During this training stage, the pretrained encoder is used for feature extraction and its weights are fixed. Stochastic Gradient Descent (SGD)[43] optimizer is used for this stage, with a learning rate of 1 *×* 10^−3^ and momentum of 9 *×* 10^−1^, which helps accelerate convergence by smoothing the trajectory of the optimization process, reducing oscillations in the gradient updates, and an exponential learning rate scheduler[44] set with gamma parameter being 9 *×* 10^−1^, which is the factor by which the learning rate is multiplied at each step.

Finally, during the last fine-tuning stage, both the autoencoder and the regressor subnet are trained simultaneously using the same parameters for the SGD optimizer and exponential learning rate scheduler, except the learning rate is reduced to 1 *×* 10^−4^. A final joint loss is used for fine-tuning the whole deep regressor. The loss *L*_*joint*_ combining *L*_*AE*_ and *L*_*R*_ is used for fine-tuning at this stage, with *λ* = 1.0 by default.

### Temporal trajectory reconstruction evaluation

To evaluate the performance of the deep regressor’s prediction, i.e., the similarity between the ground truth gene expression vector of all genes at non-selected time points **G**_**t**∉**S**_ and the predicted vector 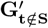, for any specific time point selection **S**, the average mean absolute error (MAE) and *R*^2^ score are computed. Among all possible selections **S** of the same number of time points |**S**| = *C*, the best time point selection **S**_**B,C**_ is defined as the one with minimal MAE:

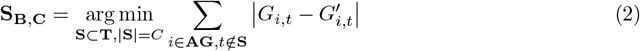

The **S**_**B,C**_ at the elbow point, denoted as **S**_**B**_, is used for subsequent single-cell inference. The selected time points from **S**_**B**_ are used as single-cell reference samples. For Fig. 2, we also computed the average MAE and *R*^2^ scores for all possible **S** with the same number of selected time points.

### Validating predicted temporal gene expression data with downstream transcriptomic analysis

To further validate DTPSP’s time-series gene expression prediction, we compare the downstream transcriptomic analysis results between the real and predicted datasets based on the optimal time point section indicated by the model.

For validating the predicted dataset’s reliability for temporal marker analysis, we identified temporal markers, i.e. DEGs between the first two time points using the real dataset, and meanwhile generate another version by substituting the second time point’s gene expression with the predicted value. For Lung Alveoli dataset and LungMap dataset with no biological replicates, genes with highest log fold changes indicated by python package diffxpy are regarded as DEGs. For the iMGL dataset with 3 replicates, standard DEG identification procedure of python package PyDESeq2[45] using t-test is used for finding the DEGs. DEGs are ranked by lowest p-values for subsequent analysis.

In computing the Hypergeometric test p-value for measuring the overlap between the two sets (ground truth and predicted) of top DEGs, the number of successes is the number of overlapping genes, the sample size and the number of successes are the number of top DEGs in the two sets and are both 100, and the population size is the number of highly variable genes involved in the analysis.

Pearson correlation analysis are performed using Python package SciPy[46]. The enrichment analysis is performed using Python package GSEAPY[47]. The top 10 enriched terms were chosen based on the highest significance (lowest p-values). After finding the top 10 terms of each, a Hypergeometric p-value was calculated for quantifying the overlap: the number of successes is the number of overlapping terms, the sample size and the number of successes are the size of two versions of compared top terms and are both 10, and population size is the total number of terms available in the Enrichr library[48]. The RRHO, shown in Fig.3b, provide an intuitive visualization of overlap between ranked gene lists by highlighting areas of significant enrichment or depletion. Each quadrant of the plot represents a specific comparison between positive or negative marker lists, with entries corresponding to the significance of overlaps based on hypergeometric tests. This approach emphasizes regions of strong statistical significance, offering insights into robust patterns of overlap. We implemented the RRHO using Python.

### Use deep generative learning to reconstruct single-cell temporal trajectory

Based on the optimal time points **S**_**B**_ selected by DTPSP’s deep regressor, we further reconstruct temporal single-cell data for the time-series study based on our deep generative learning component. The deep generative learning model is a Variational Autoencoder - Generative Adversarial Network (VAE-GAN) adapted from scSemiProfiler. It uses the single-cell data from the selected time points as reference single-cell data and decompose the other non-selected time points’ bulk data into single-cell data. After this single-cell inference process, all time points will have single-cell data available. Subsequently, DTPSP’s single-cell temporal reconstruction performance is further evaluated by comparing the inferred single-cell data and the ground truth single-cell data. In addition, we also benchmarked with scSemiProfiler with default setting, which selects reference time points by clustering the samples and selecting the samples (time points) closest to the cluster centroids as the reference.

### The architecture of the deep generative learning component for single-cell inference

The deep generative learning model employs a Variational Autoencoder (VAE) as the generator and a Multi-Layer Perceptron (MLP) as the discriminator, implemented using PyTorch based on SCVI[49]. A Graph Convolutional Network (GCN) layer was included at the beginning of the encoder via precomputing neighbor-aggregated cell data, leveraging the K nearest neighbors (K=15 by default) identified using Scikit-learn. The VAE’s encoder and decoder feature linear layers with ReLU activations, with a latent space dimension of 32 and a hidden layer size of 256. The discriminator consists of a 4-layer MLP with Leaky ReLU activations, culminating in a Sigmoid output for prediction probabilities.

### Deep generative model training

In the first pretraining stage, we train a variational autoencoder (VAE)-based generator on cells sampled from the selected time points **S**_**B**_, which constitute the most informative subset identified by our model. This generator aims to accurately reconstruct each cell’s gene expression profile from the latent space, while also countering the discriminator via an adversarial loss. Formally, the generator’s pretraining loss is defined as:

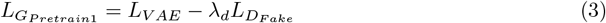

Here, *L*_*V AE*_ measures the reconstruction quality of the VAE on the reference cells from **S**_**B**_:

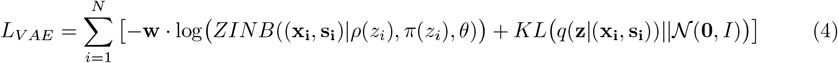

where *N* is the number of single cells from a reference sample at one of the selected time points. The vector **x**_**i**_ represents the expression values of the *i*-th cell, and **s**_**i**_ is the corresponding gene set score computed by averaging the gene expression value in the gene set. The encoder *MLP*_*Encoder*_ (*GCN* ((**x**_**i**_, **s**_**i**_))) parameterizes the latent variable **z**_**i**_ and the decoder networks *MLP*_*Decoder,ρ*_ and *MLP*_*Decoder,π*_ produce the ZINB parameters *ρ*(*z*_*i*_) and *π*(*z*_*i*_). The parameter vector *θ* is learned during training. Following this VAE pretraining, the deep regressor leverages the learned representations to predict gene expression at other time points beyond **S**_**B**_.

The adversarial component 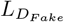 encourages the generator to produce more realistic reconstructions:

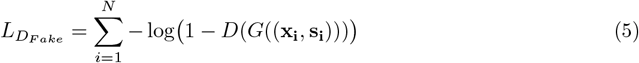

where *G*((**x**_**i**_, **s**_**i**_)) is the reconstructed cell from the generator, and *D*(*·*) is the discriminator’s predicted probability of being real. *λ*_*d*_ is a scaling factor.

The discriminator is trained to distinguish real cells from reconstructed ones:

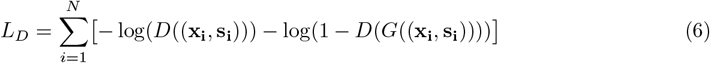

We first train the generator until convergence and then alternate between training the generator and discriminator every 3 epochs.

Subsequent to the pretrain stage, the model is further trained with a slightly modified reconstruction loss. The second training stage uses a full-batch approach without freezing any subnetworks. We incorporate an additional reference sample bulk regularization term *L*_*BulkR*_ into the generator’s loss:

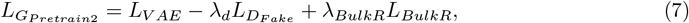

where

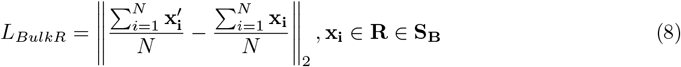

Here, **x**_**i**_ and 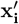 represent real and reconstructed cells from a reference time point sample **R** in the best selection **S**_**B**_, respectively. By aligning the pseudobulk signals of the generated cells with the actual pseudobulk, the model becomes better primed to integrate bulk data information into single-cell reconstructions.

Finally, the last stage, a fine-tuning stage is crucial for inferring the single-cell data the non-selected time points that are not in **S**_**B**_. In the fine-tuning phase, we discard the discriminator and focus solely on the generator’s adaptation to the target samples. Instead of using real bulk measurements directly, we rely on pseudobulk estimates to account for potential discrepancies introduced by sequencing technologies and dropout effects in single-cell data. This alignment is achieved by refining the generator with:

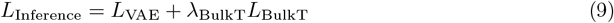

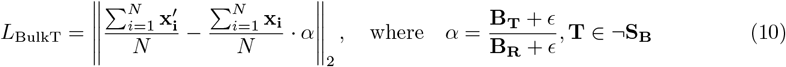

In these equations, **B**_**R**_ and **B**_**T**_ denote the pseudobulk profiles of the reference time point sample from **S**_**B**_ and a target sample **T** not in **S**_**B**_, respectively, and *α* is the scaling factor used to guide the generator in producing single-cell expressions for the target samples. This fine-tuning step ensures that the model can produce realistic single-cell profiles that accurately reflect the target’s underlying bulk expression patterns, thereby enabling effective *in silico* inference of single-cell data at minimal cost.

### Single-cell inference performance evaluation

To quantitatively evaluate inference accuracy, we adopted the error metric from scSemiProfiler to assess the fidelity of semi-profiled samples compared to their real-profiled counterparts. The evaluation process involves the following key steps:

#### Data aggregation and dimensionality reduction

For error computation and visualization to compare real-profiled (ground truth) and semi-profiled datasets, all samples (real-profiled or inferred using the deep generative model) from the same study are combined into a single dataset before dimensionality reduction. The term “semi-profiling” refers to generating single-cell data for an entire dataset, either through direct sequencing of selected reference samples or in silico inference using the deep generative model. Principal Component Analysis (PCA) [50, 51, 52] is then applied to this combined dataset to reduce its dimensionality. Specifically, we retain *D* = 100 principal components to capture the most significant variance in the data. Let 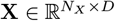 represent the PCA-transformed real-profiled data of a sample, and 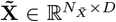 denote the corresponding inferred single-cell data, where *N*_*X*_ and 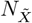 are the numbers of cells in the real and semi-profiled datasets, respectively.

#### Nearest neighbor identification

For each cell **x**_*i*_ in **X**, we identify its *K* = 1 nearest neighbor in 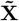, and conversely, for each cell 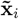 in 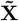, we identify its *K* = 1 nearest neighbor in **X**. We utilize the FAISS library [53] for efficient nearest neighbor search.

#### Single-cell level difference metric

The single-cell level difference metric, denoted as Dif_sc_ 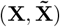, is defined as:

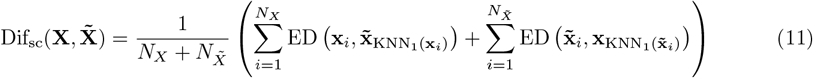

where KNN_1_(**x**_*i*_) denotes the index of the single nearest neighbor of cell **x**_*i*_ in 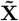, KNN_1_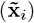 denotes the index of the single nearest neighbor of cell 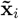 in **X**, and ED( *·,·*) represents the Euclidean distance between two cells.

#### Aggregate single-cell inference error

The average inference error, 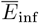, is calculated as the mean of Dif_sc_ 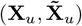 across all *U* samples: 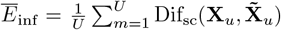 where, *U* is the total number of samples, Dif_sc_ 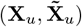 is the single-cell difference metric for the *u*-th sample.

#### Global bound computation

To contextualize the inference error, we compute global lower and upper bounds using the aggregated data: **Lower bound (LB):** The combined dataset is randomly shuffled and split into two equal halves. The lower bound is then calculated as the mean Euclidean distance between each point in the first half and its nearest neighbor in the second half, averaged over both halves.**Upper bound (UB):** The upper bound is determined by pairing each real-profiled sample with a randomly selected inferred sample and computing the mean Euclidean distance between these pairs. These bounds are computed once per round across all projects to ensure consistency in error evaluation.

#### Error normalization

Each sample’s error is normalized using the computed bounds to facilitate consistent comparisons across multiple runs and conditions: 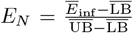, where 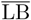 is the mean lower bound, 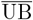 is the mean upper bound. This normalization scales the error metrics to a standardized range, enabling scalable and comparable performance evaluations across different experimental setups.

### Single-cell analysis for downstream tasks evaluation

#### Cell type annotation

The annotation of the inferred single-cell data is performed using a multilayer perceptron (MLP) classifier in Python package scikit-learn trained on the annotated reference single-cell samples. For iMGL and LungMap datasets, we retained their original processed annotations, removing cells lacking confident annotations. For the Lung Alveoli dataset, the original cell type annotation is not available. So we manually annotated the cell types by first performing PCA transformation, computing neighbor graph, applying Leiden clustering, all implemented in SCANPY, and then label each cluster based on marker gene expression. Clusters with the same cell types are merged and cell types represented by fewer than 50 cells are removed to reduce noise and enhance the robustness of subsequent analyses. The PCA, neighbor graph construction, and UMAP [54] visualization are all based on SCANPY’s implementations.

#### Dimensionality reduction and visualization

To visually assess the quality of inferred single-cell data, we merged both the real-profiled and semi-profiled datasets into a single expression matrix. Principal Component Analysis (PCA)[50, 51, 52] was then applied to extract the primary axes of variation using 100 principal components. Utilizing these principal components, we constructed a K-nearest neighbor (KNN) graph with 50 neighbors to capture local cell relationships and employed Uniform Manifold Approximation and Projection (UMAP) [54] to project the data into two dimensions. This joint embedding allowed for direct comparison between inference and real-profiled cells, offering a straightforward qualitative assessment of how closely the inferred data resembled the authentic cellular landscapes.

#### Enrichment analysis

To confirm that inferred data preserved meaningful biological signals, we identified the top 100 differentially expressed genes (DEGs) characterizing each cell type in both the real-profiled and semi-profiled datasets. Using these DEGs, we performed GO or Reactome pathway enrichment analysis via GSEApy [47]. We compared the significance and overlap of enriched terms between real and semi-profiled data to assess fidelity in capturing functional insights. A hypergeometric test was used to evaluate the overlap in identified DEGs, while Pearson’s correlation (computed using SciPy [46]) measured the similarity in enrichment score distributions. This ensured that the semi-profiled data not only approximated the cell-level gene expression patterns but also retained underlying biological context.

## Data availability

Three publically available datasets are used to evaluating our model. The Lung Alveoli data is from Zepp et al.’s study[33] and can be downloaded from the Gene Expression Omnibus (GEO)[55] repository under accession number GSE149563. The LungMap dataset was taken from the LungMap Consortium within the Single-cell RNA-seq section on September 1st, 2024, downloading data with LungMAP IDs LMEX0000003683 till LMEX0000003689. Finally, the iMGL dataset is from Ramaswami et al.’s study and can be downloaded from the GEO repository under accession number GSE226081.

## Code availability

The source code for DTPSP is publicly available on GitHub at https://github.com/mcgilldinglab/DTPSP.

## Acknowledgments

This work was funded in part by grants awarded to [JD]. We gratefully acknowledge the support from the Canadian Institutes of Health Research (CIHR) under Grant Nos. PJT-180505; the Funds de recherche du Québec - Santé (FRQS) under Grant Nos. 295298 and 295299; the Natural Sciences and Engineering Research Council of Canada (NSERC) under Grant No. RGPIN2022-04399; and the Meakins-Christie Chair in Respiratory Research.

## Notes

https://www.ncbi.nlm.nih.gov/geo/query/acc.cgi?acc=GSE149563

https://www.lungmap.net/

https://www.ncbi.nlm.nih.gov/geo/query/acc.cgi?acc=GSE226081

### Competing Interest Statement

The authors have declared no competing interest.

