## Supplementary Figures for "DTPSP: A Deep Learning Framework for Optimized Time Point Selection in Time-Series Single-Cell Studies"

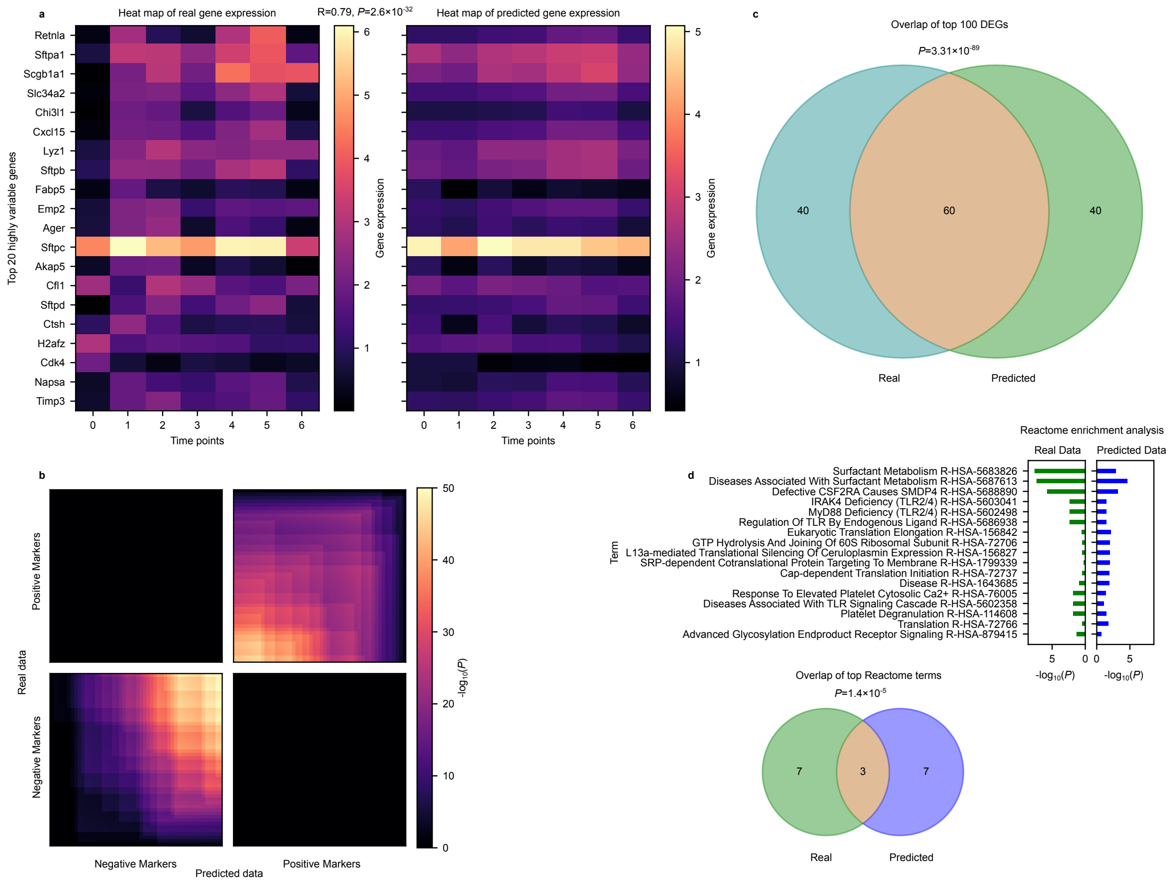


**Fig. S1**: **Biological validation of DTPSP predictions on the LungMap dataset. a,** Heatmap comparing the expression pattern of top 20 differentially expressed genes (DEGs). DEGs' expression in the predicted dataset (right heatmap) is highly similar to the expressions in the predicted dataset (left), justified by a Pearson correlation of ($R = 0.79$) and p-value ($P = 2.60 \times 10^{-32}$) between the two flattened matrices. **b,** Rank-Rank Hypergeometric Overlap (RRHO) plot comparing positive and negative marker genes between the real and predicted LungMap datasets. Bright regions in the first and third quadrants indicate significant concordance of markers, as reflected by hypergeometric p-values in these regions. **c,** Venn diagram illustrating the overlap of the top 100 differentially expressed genes (DEGs) identified from the real and predicted LungMap datasets. A substantial overlap of 60 genes is observed, supported by a hypergeometric p-value of $3.31 \times 10^{-89}$, demonstrating the reliability of DTPSP in recovering biologically relevant DEGs. **d,** GO enrichment analysis comparing the real and predicted LungMap datasets. The union of the top 10 enriched Gene Ontology (GO) terms is shown, with a Venn diagram highlighting a statistical significant overlap.

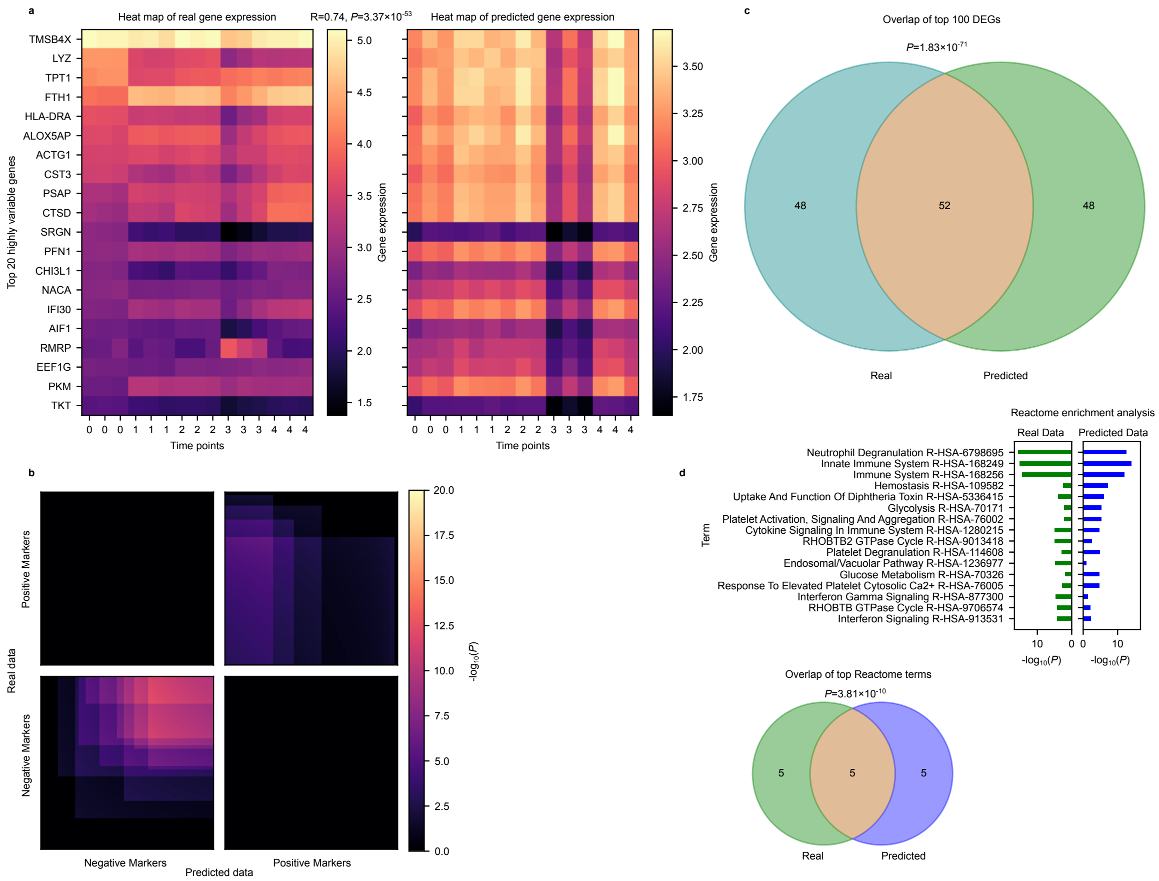


**Fig. S2**: **Biological validation of DTPSP predictions on the iMGL dataset with real bulk data. a,** Heatmap comparing real and predicted gene expressions. The top 20 DEG's expression patterns in the two replicates of real dataset and predicted dataset are shown. Pearson correlation 0.74 and p-value $3.37 \times 10^{-53}$ demonstrate a strong match between real and predicted data. **b,** Rank-Rank Hypergeometric Overlap (RRHO) plot showing the similarity of positive and negative marker genes. Bright regions in the first and third quadrants highlight areas of significant overlap. **c,** Venn diagram illustrating the overlap of the top 100 DEGs. 52 overlapping genes confirm the method's ability to identify biologically relevant DEGs. **d,** GO enrichment analysis comparing the top 10 enriched Gene Ontology (GO) terms. The Venn diagram shows 5 shared terms, supporting the consistency of DTPSP's pathway predictions.
